# Darwin’s “neuters” and the evolution of the sex continuum in a superorganism

**DOI:** 10.1101/2023.11.13.566820

**Authors:** J Oettler, T Wallner, B Dofka, J Heinze, N Eichner, G Meister, M Errbii, M Rehli, C Gebhard, E Schultner

## Abstract

Ant castes are an amazing example of phenotypic plasticity. In worker-destined embryos of the ant *Cardiocondyla obscurior*, the default female developmental trajectory is interrupted even before the gonadal precursor cells acquire a sexual identity. miRNA and mRNA expression in embryos reveal three distinct phenotypic entities: males, females, and “neuters”, as Darwin coined the worker caste in “On the Origin of Species”. Based on these results we propose that haplodiploidy, in conjunction with insect sex determination, allows for the expression of a third dimension on the sex continuum, thus facilitating the evolution of individuals which develop traits their parents do not have.

**One sentence summary:** Sex and caste differentiation begin during the same embryonic developmental window in the ant *Cardiocondyla obscurior*.

## Main text

How ant caste development works and how it evolved are outstanding and unsolved questions (*1*, *2*). Early work described the presence of wing discs in worker larvae (*3*), thereby showing that worker ants, though wingless as adults, retain parts of the queen developmental body plan. Although few data were available, ant caste determination quickly became a debate over “blastogenesis” (which unlike the strict definition of the term meant to express that caste was determined genetically) versus “trophogenesis”, meaning determination via nutritional factors during larval development (*4–6*). Today, we know that, depending on the species, genetic, maternal and environmental factors can all be involved in queen-worker caste determination (reviewed in (*7–9*)). Still, the view persists that nutritional variation during larval development, and its effects on body size, regulate caste determination and differentiation across the ant phylogeny (*10–12*) analogous to insights from the honey bee (reviewed in (*13*)) and studies of minor and major ant worker castes (*7*, *14–16*).

The idea that ant caste differentiation only occurs during larval development ignores that the defining difference between ant queens and workers lies in their reproductive organs, the formation of which begins during embryogenesis. In female insects, ovaries develop from mesodermal somatic gonadal cells incorporating germ cells under the influence of the endoderm (*17*), whereas the spermatheca, the sperm storage organ of insects, is an ectodermal structure originating from cell clusters specified in the embryo (*18*). Despite the different germ layer origin, both ovaries and spermatheca are typically constrained in their growth and development in ant workers (*19*). In few ant species, workers grow a fully functional queen-like reproductive apparatus, including ovaries and a spermatheca (*18*, *20*, *21*). In most species, workers exhibit rudiments of the spermatheca or have lost the organ altogether but develop ovaries - generally with fewer ovarioles than the queen - which can produce eggs that give rise to haploid males or serve a trophic function. Workers completely lacking sex cells and reproductive organs, i.e., without the ability to produce gametes, have evolved convergently in ca. 3% of the total ∼300 ant genera (in 2 of the 16 extant subfamilies). Gene expression studies in two species with sterile workers –

*Cardiocondyla obscurior* and *Monomorium pharaonis* - show that developmental trajectories of queens and workers are distinct in late larval stages (*22–24*). Earlier divergence has been proposed to occur in *M. pharaonis*, where germ cells in some embryos appear to degenerate shortly after the germ line has been established, indicating an early embryonic point-of-no-return to the worker trajectory (*21*). However, *M. pharaonis* embryos do not show caste-specific patterns of gene expression (*24*), contrasting the tenet of eco-evo-devo that embryonic development underlies the evolution of major traits such as sterility.

The two-colored heart-node ant *Cardiocondyla obscurior* is a laboratory model for the study of social evolution with workers which entirely lack reproductive organs (*25*). The discovery of queen control over caste allocation (*26*), and of a character that allows non-invasive identification of queens from late embryogenesis onwards (*27*) point to an embryonic segregation of the developmental pathways of queens and workers in this species. Here, we reconstructed embryo- and gonadogenesis and compared RNA expression of queen, male and worker late-stage embryos to demonstrate that both sex and caste differentiation begin during the same embryonic developmental window.

## Results and Discussion

### Ovarian development

Gonadogenesis in ants has not been described, so we used development in *Drosophila melanogaster* as a guide. Here, gonadogenesis begins with primordial germ cells (PGCs), which form from maternally provided cytoplasm at the posterior pole of the embryo and migrate towards anterior with germ band elongation to enter the embryo with the midgut invagination step. The parasegments 10-12 initiate differentiation of somatic gonadal precursor clusters (SGPs) involving the homeotic genes *abdA* and *AbdB*, which signal their location via AbdB to the “waiting” PGCs in the midgut. The PGCs then migrate through the midgut epithelium to posterior to merge with the SGPs, together giving rise to the gonads (*28*, *29*). The coalesced gonadal precursors also express *nanos*, required for germ cell specification (*30*, *31*).

In *C. obscurior*, embryonic development lasts for approximately nine days, and we identified five main stages (S1-S5, Supp Fig S1). Early-stage embryos express *nanos* at the posterior end (Supp Fig 2). In late-stage S3 embryos, 5-6 days after egg-laying (AEL), when germ band retraction is completed, the PGCs have migrated posterior and coalesced into three cell clusters with the SGPs (Fig 1a, Supp Fig 2). The gonads are visible as two posterior bilateral clusters that co-express *AbdB* and *nanos* in S5 embryos shorty before hatching (Fig 1b).

**Fig 1.**
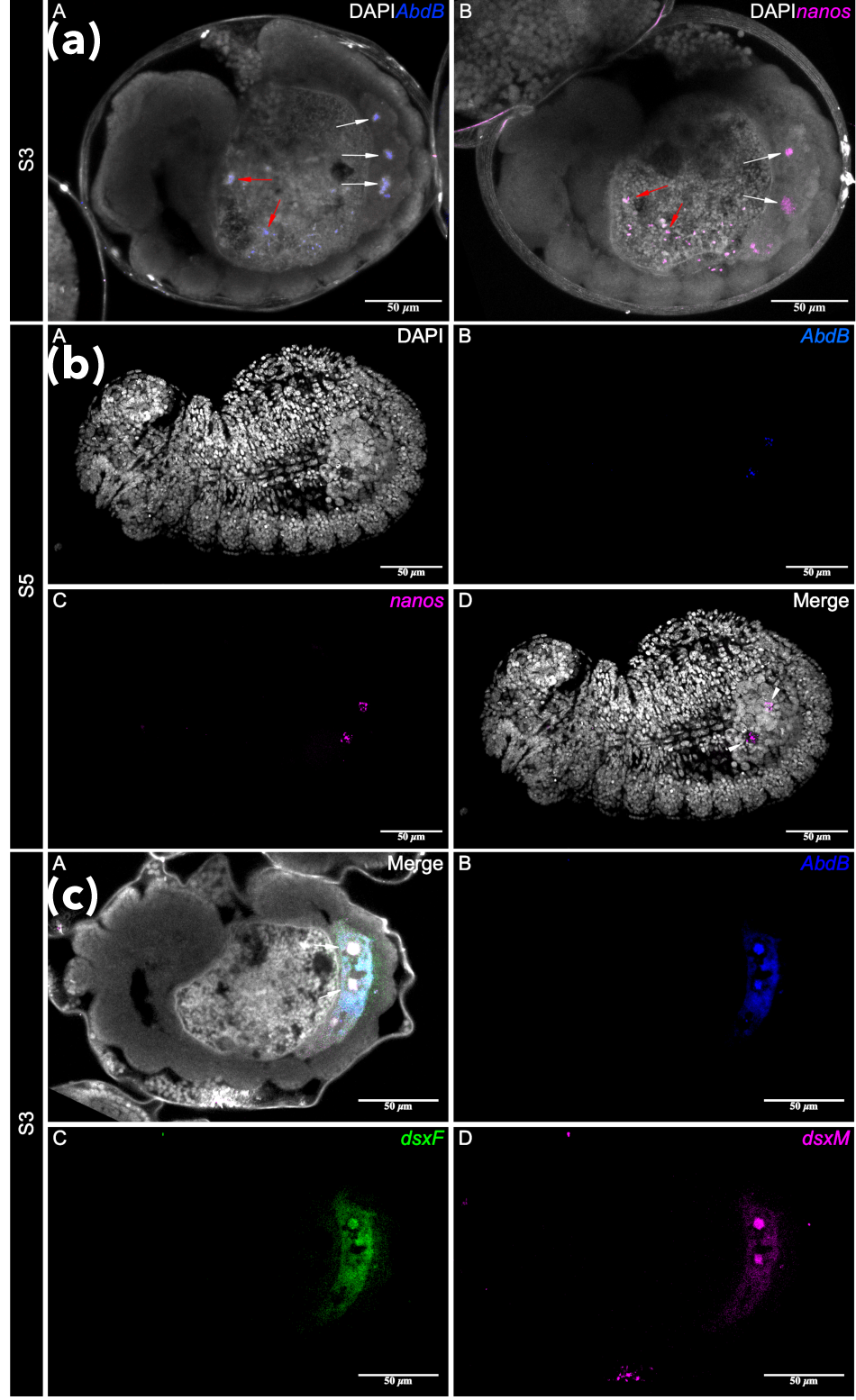
**(a) *AbdB* is required for specification of somatic gonadal precursor clusters (SGPs) in S3 embryos.** (A) *AbdB* is expressed in cluster of cells resembling three parasegments (white arrows), indicating the presence of SGPs. Cells in the midgut also express *AbdB* (red arrows) (B) Germ cells expressing *nanos* and *AbdB* are found in the same parasegments coalescing with the SGPs (white arrows). Cells in the midgut also express *nanos* (red arrows). Grey = DAPI. **(b)** Gonad formation in S5 embryos. (A) Stage 5 embryo counterstained with DAPI against cell nuclei. (B) Expression of *AbdB* marking SGPs. (C) Germ cells expressing *nanos*. (D) The embryonic gonads coalesce into two cell clusters co-expressing *AbdB* and *nanos* (arrows). **(c)** Co-expression of *dsxF* and *dsxM* isoforms in SGPs in S3 embryos. (A) Stage 3 embryo co-expressing the known *doublesex* isoforms *dsxF* and *dsxM* in SGPs (arrows). (B) *AbdB* expression marking SGPs. (C) Expression of *dsxF*. (D) Expression of *dsxM*.

**Fig 2.**
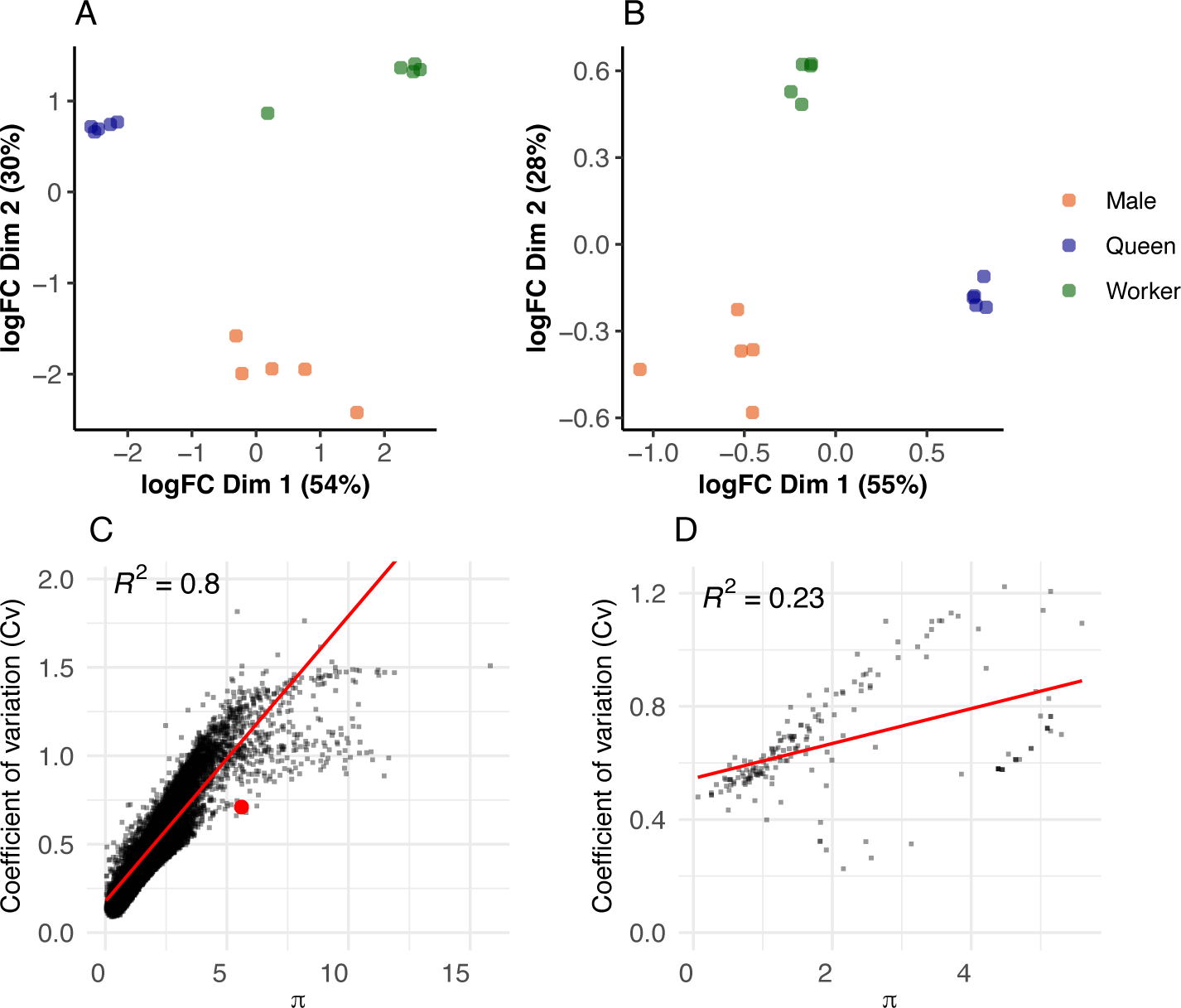
A) MDS plot of mRNA expression in *C. obscurior* queen, worker and male embryos. B) MDS plot of the miRNA expression in *C. obscurior* queen, worker and male embryos. C) Correlation of the plasticity index π with the coefficient of variation C_V_ for each mRNA, highlighted in red is *dsx*. D) Correlation of the plasticity index π with coefficient of variation C_V_ for each miRNA.

### Sex and caste differentiation co-occur during early gonadogenesis

There are no known sex- or caste-specific genes or chromosomes in ants, including our model *C. obscurior* (*22*, *32*), so that sex- and caste-specific growth and development must result from differential expression of the same genes. A link between sex and caste determination and differentiation in ants has been proposed (*23*, *33*, *34*) but the proximate mechanisms and evolutionary processes underlying this presumed co-option are largely unclear.

The insect *doublesex* (*dsx*) gene functions as a molecular switch, with alternative splicing specifying sexual identity (*35*). In *D. melanogaster*, the body is essentially a mosaic of tissues that express *dsx* isoforms (the gonads and e.g. regions of the CNS), and tissues that do not express *dsx* and thus have no sex-linked phenotype (*36*). In *C. obscurior* S3 embryos, expression of the two described *dsx* (Cobs15665) isoforms *dsxF and dsxM* (*33*) is restricted to the abdominal regions containing the precursors of the embryonic gonad, the SGPs (Fig 1c). This indicates that in S3 embryos, the sexual identity of the SGPs is determined and that expression of both isoforms is necessary for female gonadogenesis.

Not all S3 embryos showed signs of *nanos*, *dsx* or *AbdB* expression, indicating these were worker embryos in which PGCs were aborted earlier. Absence of evidence is not evidence of absence, especially given that ant embryos are difficult to stain. But this tentatively suggests that in *C. obscurior*, the worker caste is determined even before the the PGCs and SGPs coalesce into the developing gonads.

### Queen, worker and male embryos differ in mRNA and miRNA expression

During the final stage before hatching (S5, 8-9 AEL) the head changes its orientation from ventral to anterior-ventral, and the embryo begins muscular movement. The female gonads begin to develop and associate with distinct crystalline deposits (of unknown cause and function), which make it possible to identify queen and worker embryos with ∼95% and ∼80% precision, respectively (*27*). To better understand embryonic caste differentiation, we sequenced mRNA and miRNA of S5 queen, worker, and male embryos (n=5 pools of 100 embryos each. mRNA: 70bp single-end, Illumina Nextseq 500/550, 31.29±2.5 SD million reads per sample, of which 9.71±1.78 million reads mapped to exons. miRNA: 80bp single-end, Illumina MiSeq, ∼1.93 ± 0.39 million reads per sample).

Of the 10113 expressed genes in the data, 8108 genes (80.2%) differed between groups (limma-voom, adjusted p < 0.01, Online Supp Table 1). This is a higher proportion than in a previous RNAseq study comparing individual third instar larvae (*22*), however these studies are not directly comparable. In addition to different methods used, pooling increases the abundance of rare transcripts, normalizes environmentally-induced individual variation, and reduces the effect of misidentified individuals.

A multidimensional scaling (MDS) plot revealed three discrete groups (Fig 2A). No caste or morph stands out prominently, instead Dim1 separates workers from queens with males in between, and Dim2 is driven by queen- and worker-biased expression. A Gene Ontology analysis showed that a larger number of GO terms were specific for the queen/worker comparison (39/48 terms, 81.3%; Online Supp Table 1), compared to the queen/male (22/40, 55%; Online Supp Table 1) and worker/male (20/35, 57.1%; Online Supp Table 1) comparisons. Also, male/queen and male/worker comparisons shared more GO terms (11), than they shared with the queen/worker comparison (5 and 3, respectively). Together, this indicates that in addition to many shared GO terms, more unique processes are involved with queen/worker differences compared to the contrasts involving males.

Of the 358 miRNAs predicted by miRDeep2 (Online Supp Table 2), 136 (37.9%) were differentially expressed between groups (limma-voom, p < 0.001). As with mRNAs, miRNA expression in worker embryos was equally distinct from that in queens and males (Fig 2B). That Dim1 is driven by queen-biased expression points to the possibility that miRNAs involved with queen/worker differences were co-opted from an ancestral sex differentiation (queen/male) mechanism.

To study the behavior of mRNAs and miRNAs which play a significant role in phenotypic plasticity in this highly variable dataset, for each gene we calculated the plasticity index (π), a measure for summarizing the versatility and expression bias of a given gene across multiple groups (*32*), and the coefficient of variation (C_V_) of that gene across all samples. In a study comparing RNAseq data of third instar larvae, the correlation between π and C_V_ was positive and highly significant (*32*), suggesting that genes with a high π and correlated with differences between phenotypes (with ’anticipatory’ or ’adaptive’ plasticity) tend to be more responsive to the environment (showing ’passive’ plasticity), compared to constitutively expressed genes.

Similarly, in the embryo mRNA data the overall correlation between π and C_V_ was strong (Fig 2C, adjusted R-squared: 0.7996, F-statistic: 4.034e+04 on 1 and 10111 DF, p < 0.001). As expected for a conserved major effect transcription factor, *dsx* (*Cobs15665*) exhibited a high π, being weakly expressed in males but showing a 14.3-fold and 18.8-fold expression increase in queens and workers, respectively (Online Supp Table 1, Supp Fig 3, p-values of pairwise Mann-Whitney-U-tests with Bonferroni-Holms p-value correction: M/Q = 0.012, M/W = 0.012, Q/W = 0.056). *dsx* exhibited a moderate C_V_, suggesting that its overall expression is under relatively tight control (Fig 2C). High levels of *dsx* in workers highlight the importance of this gene in regulating worker differentiation during late embryogenesis - keeping in mind that workers lack reproductive organs and sex cells at this stage.

The positive correlation between π and C_V_ was also highly significant in the miRNA data, but less strong due to several miRNAs with exceptionally high π (Fig 2D, Adjusted R-squared: 0.2284, F-statistic: 55.17 on 1 and 182 DF, p < 0.001), which may be involved with the regulation of phenotypic divergence. These need to be validated in the lab before candidate genes can be identified and tested for possible interactions with e.g. *dsx*.

Together the expression data 1) confirm that caste is fixed by the final embryonic stage in *C. obscurior*; 2) show that post-transcriptional regulation via miRNAs plays a compelling role in caste and in sex differentiation; 3) demonstrate that miRNA expression is under tighter regulation than mRNA expression, as indicated by lower C_V_ values; this is in line with the role of miRNAs in post-transcriptional regulation; 4) reveal a number of mRNA and miRNA candidates with high π and low C_V_ which we predict to be associated with processes pertaining to developmental plasticity, and that are of particular interest for future bioinformatic and functional evo-devo studies.

## Conclusion

A major realization to the aspiring myrmecologist is that ant castes presented a serious challenge to Charles Darwin. Not because it was difficult for him to envision an altruistic worker caste; indeed, years later inclusive fitness theory explained how worker sterility can evolve under kin selection (*37*, *38*). Instead, it was the development of offspring with morphological traits distinct from those of their parents, and in the absence of genetic differences, which seemed difficult to explain with a theory of natural selection (*39*). Today, it is established that developmental plasticity allows the expression of distinct phenotypes from the same genetic information (*40*). Studies comparing the development of reproductive organs in ant species spanning the range of worker reproductive constraints have revealed different timepoints when development of reproductive organs can be interrupted, and that these correspond to the state of spermatheca and ovary development in adult workers (*18*, *19*, *21*). Studies on the growth and development of wing discs have likewise revealed different interruption points (*41–46*). Finally, caste-specific growth and development during larval stages are reflected in differential gene expression (*22–24*, *32*). Together, these results have spurred a discussion about the evolution of ant caste development (*1*, *2*, *10*, *11*), bringing new attention to the century-old question of how ant queens and workers are made.

Here, we show that caste- and sex-specific growth in *C. obscurior* is initiated during embryonic development, and that late-stage embryos of workers, queens and males are distinct in both mRNA and miRNA expression. Based on these results, we propose a possible solution for how ants achieve extreme levels of phenotypic plasticity grounded in the principles of haplodiploidy and insect sex determination. Chromosomal sex determination systems such as XX/XY, XX/X0, and ZW/WW usually have a dichotomous female or male outcome, encoded by genes located on sex-specific chromosomes. Many species deviate from the norm and exhibit adaptive variations, ranging from androgenesis and gynogenesis to hermaphroditism of various forms, the elimination of males in parthenogenetic species, or the elimination of pure females in species with androdioecy. Nevertheless, in general, individuals inherit or acquire one sex and/or the other to produce gametes. In contrast, in haplodiploid insect species, the difference between the sexes lies only in the number of chromosomes so that phenotypic variation is solely based on expression differences of the same set of genes. Thus, a haplodiploid species is not constrained to only two sexual identities.

This is evident in *doublesex* expression, a transcription factor with conserved function in sexual development in insects. In *Drosophila dsx* acts as a transcription factor that generates sex-specific isoforms to regulate sex-specific development. Production of the female- specific *dsx* isoform requires the presence of functional Sex-lethal (Sxl) protein, which is expressed exclusively in female flies that contain two X chromosomes. Sxl affects *dsx* production via the regulation of alternative splicing of transformer (*tra*), enabling the production of functional tra protein, that, together with tra2, governs female-specific splicing of *dsx*. Ants have two *tra* copies (*47*), whose functions are unknown, and an ortholog of *sxl*, but no sex chromosome. Workers in *C. obscurior* differ from queens and males in the expression of the female and male *dsx* isoforms across different developmental stages (third instar larvae, pupae, adults) demonstrating that there are more than two *dsx* expression patterns (*33*). Analogous, queen SGPs differentiate under the influence of the two known *dsx* isoforms, whereas workers, lacking gonads, do not exhibit *dsx* specification of SGPs in S3 embryos. Foregoing the default female sexual identity at an early embryonic stage allows for later expression of the same set of developmental genes in a third phenotypic context, which is not male or female, but “neuter”. In support of this idea, shortly after the gonads have formed and acquired a sexual identity in queens and males, late-stage worker embryos differ as strongly in mRNA and miRNA expression from queens as they do from males. Many processes appear to be specific for the queen/worker differentiation, including *dsx*, which is more highly expressed in queens compared to males, but even more so in workers. Furthermore, it appears as if miRNAs involved with sexual differentiation also play a role in the development of workers. Together with correlative data showing that sex and caste differentiation during larval development are linked (*23*, *33*, *34*), our results provide evidence for a co-option of sex differentiation pathways in ant caste development.

For caste differentiation to occur during embryogenesis, it must be preceded by caste determination. In *M. pharaonis*, it was proposed that worker caste determination in the embryo results from the degeneration of germ cells (although the stainings could also show migrating PGCs) (*21*). We think that worker sterility in *C. obscurior* may be initiated even earlier by an inhibition of PGC specification through interfering maternal epigenetic factors, as queens seem to have full control over caste determination (*27*). Physiology probably plays a decisive role as a Zeitgeber in this process since queens increase production of queen-destined eggs with age (*26*). Perhaps this employs growth hormones and other maternally derived factors such as RNAs and proteins, as has been suggested in temperate ants with maternal egg-biasing (*48–50*); these may be transferred to oocytes via extracellular vesicles or via vitellogenin (*51*). The occurrence of maternal egg biasing even in species in which workers retain functional ovaries (*50*, *52*, *53*) suggests that maternal effects on ant caste may be more widespread. Perhaps it is time that the debate over nature vs. nurture in ant caste development (*12*) shifts focus, asking instead what forms of maternal provisioning have evolved across the ant phylogeny and how these interact with embryonic development.

## Methods

### Species

*Cardiocondyla obscurior* is a minute myrmicine ant (queens 3mm, workers 2mm) distributed around the globe in warm climates, with small colonies that live in cavities in trees and shrubs, e.g., under bark, in knotholes, rolled leaves, and aborted fruits (Supp Fig 4). The genus is characterized by the evolution of wingless fighter males in addition to the ancestral winged males, and incestuous, sometimes even oedipal mating to produce diploid female offspring from small propagules, thus facilitating a quasi-clonal tramp lifestyle (*54*, *55*). Workers have no wing imaginal discs in third instar larvae while wingless males have remnants of wing discs, suggesting an early to mid-larval developmental switch between winged and wingless males (*42*), reminiscent of worker caste determination in *Pheidole* ants (*14*). *C. obscurior* has the smallest known ant genome (∼193MB (*56*) and is associated with an enigmatic endosymbiont, *Cand.* Westeberhardia, also with a highly eroded genome (∼530kb (*57*, *58*). Development takes ca. 30 days from egg laying to adult, queens mate within the first two weeks of their lives, start egg-laying within a further two weeks, and have the same pace (lifespan ∼28 weeks) and shape of aging as workers (*26*, *59*). Queens can be identified from late embryogenesis onwards by the presence of paired crystalline deposits associated with the developing ovaries (*27*).

The ants used here are of the ancestral Asian lineage (’Oyp B’, Okinawa, Japan), and have been in the laboratory since 2011 (>70 generations) (*60*). Colonies were kept in square plaster-bottom nests (100 mm x 100 mm x 20 mm, Sarstedt, Germany) with plastic inserts containing three chambers covered in dark foil, in a climate chamber under a 12h/12h and 22°C/26°C night/day cycle at 80% humidity. Colonies were fed three times a week with honey, fruit flies and pieces of cockroaches and provided with water ad libitum.

### Embryogenesis

We followed egg development by setting up colonies with single mated queens and 10 workers from stock colonies. Queens were left to lay eggs for 24 hours and then removed thereafter. Eggs were tended to by nurse workers while being aged. Eggs were collected every 24 hours to determine the developmental stage of the embryo. The selected eggs were submerged in a dissection dish containing phosphate-buffered saline solution PBT (0.3 %) and were then transferred onto a microscope slide. The slides were sealed with nail polish and imaged using a stereomicroscope connected to a camera (Keyence VHX 500FD, Neu-Isenburg, Germany). Images were then processed using the Fiji distribution (*61*) in the open-source platform ImageJ, and the software ScientiFig (*62*).

### Whole mount in-situ hybridization

Eggs were collected from stock colonies and submerged in a dissection dish containing PBT (0.3 %). The embryos were visually classified into their respective stage, and then used for in situ hybridization. Fluorescent in situ probes were designed for the germline genes *nanos*, the female- and male specific splice forms of *doublesex* (*dsxF*/*dsxM*) and *Abdominal B* (*AbdB*) (Supp Table 3) using the Stellaris RNA FISH Probe Designer (Biosearch Technologies, Inc., Petaluma, CA). *dsxF* mRNA was labeled with Quasar 570, *dsxM* and *nanos* mRNA with Quasar 670 and *AbdB* with CAL Fluor Red 610 dye. The Stellaris protocol for *Drosophila* embryos was adapted to our ant (Suppl. Protocol 1). Images for in situ hybridization were visualized using a Zeiss LSM 880 and LSM 980 with Airyscan 2 laser scanning microscope under 40x objective lenses. Images were then processed using the Fiji distribution (*61*) in the open-source platform ImageJ, and the software ScientiFig (*62*). The appearances (’Color’) of the final Fig1(b) and Fig1(c) were adjusted automatically in Preview (macOS Ventura).

### mRNA and miRNA sequencing Sampling and sequencing

For each of the three egg phenotypes (male, worker, queen), five pools containing 100 S5 stage embryos each were sampled. To identify embryo caste, eggs were collected from the colonies and transferred into PBT (0.3%) for closer inspection under a stereomicroscope. Identified samples were immediately frozen in liquid nitrogen and stored at -80 °C until extraction of mRNA and miRNA.

To collect male eggs, 24 colonies containing virgin queens were set up, as these lay only haploid, male destined eggs. Each colony was started with three queen pupae and ten workers. Worker numbers were kept constant by replacing dead workers from the respective stock colonies once a week. Colonies were checked three times a week for the presence of eggs. Eggs from the first week were discarded since these may have been accidentally transferred while establishing the colony. After around 4 weeks all colonies had initiated egg laying. To synchronize egg laying, all eggs from all colonies were collected and discarded on the same day. After five to seven days, all eggs from the virgin colonies were collected and inspected under a stereomicroscope and all S5 embryos were sampled. After five eggs were collected, colonies were left to rear further eggs into pupae, to verify that all colonies only produced males.

Female eggs were collected from a total of 20 stock colonies. Caste was identified under a stereomicroscope based on the presence of crystalline deposits of a yet unknown substance, which appear at the beginning of the germband retraction in the S3 stage of queen, but not worker destined embryos. In 1st instar worker larvae these crystalline deposits also appear but are never paired. To prevent a collection bias, queen and worker embryo samples were always collected simultaneously. We have no way of distinguishing worker from male embryos to date and ca. 5-10% of the total brood is male (*26*), so the “worker”-pools likely contain some male embryos. “Queen” and “male” pools are pure with very high probability.

Total RNA was extracted using the Trizol (Thermo Fischer Scientific, Waltham, MA, USA) RNA extraction protocol. Eggs were crushed using a pestle for 15 seconds, before adding 500 μl of the Trizol reagent. After crushing the eggs for another 30 seconds, an additional amount of 500μl of Trizol was used to rinse remaining tissue from the pestle. After that the standard protocol was used.

The mRNA libraries were generated using the Illumina Stranded mRNA Prep kit (Illumina Inc., San Diego, CA, USA) and analyzed using Qubit HS dsDNA and TapeStation HSD1000 reagents (Agilent). RNA-seq libraries were single-end sequenced at a length of 86bp on the NextSeq 550.

From the same samples microRNA was purified using a modified version of the Small-RNA- Cloning with the TrueSeq-System (*63*). For cloning, 0.2 – 0.5μg of RNA of each sample were used. In short after initial ligation of a 3’ adapter the ligation products were purified via PAG electrophoresis. Then an RNA 5’ adapter was ligated, followed by cDNA synthesis and PCR amplification, where the functional sequence elements for Illumina sequencing were introduced. The resulting PCR amplicons were subjected to a second gel electrophoresis to purify the library amplicons representing miRNA inserts. These were eluted from the gel, precipitated and finally resolved in water. Quality assessment was performed using Tapestation (Agilent Technologies) and quantitative PCR. After equimolar pooling sequencing was performed on an Illumina MiSeq platform.

### Bioinformatics Transcriptome analyses

After assessing the raw reads quality using FastQC [https://www.bioinformatics.babraham.ac.uk/projects/fastqc/], we removed ribosomal RNA (rRNA) sequences using SortMeRNA (*64*) and the silva databases (www.arb-silva.de). The rRNA-free single-end reads were aligned to the Cobs3.1 reference genome (PRJNA934066; Errbii et al. submitted) using STAR (*65*), and mapping quality was evaluated with Qualimap (*66*). We then used FeatureCounts (*67*) to count reads per annotated gene in the reference for downstream differential gene expression analysis. On average, 97% of the reads mapped to a unique position and approximately 10 million reads per sample were used to produce the count data (Online Supp Table 1). Differential expression (DE) analysis was performed using the R package limma-voom (*68*, *69*). Read counts were converted to log2 counts per million (CPM) and genes were filtered for those with at least 20 reads. We applied the method of trimmed mean of M-values (TMM) from edgeR (*70*) to calculate normalization factors and to account for library size variations. Differentially expressed genes were identified using the voom function in the R limma package, and filtered under a significance threshold (p < 0.01) with Benjamini-Hochberg correction. Additionally, we computed plasticity indices (π) (*32*) and coefficients of variation (C_V_) for each gene to assess overall gene expression plasticity and variability, respectively. A similar approach was employed for miRNA DE analysis, filtering out miRNAs with fewer than 10 mapped reads. A summary of all computed metrics, including logarithmic fold-changes (logFC), adjusted p-values and π for genes and miRNAs can be found in Online Supp Table 4.

For Gene Ontology (GO) enrichment analyses, we used the R package topGO (*71*) to identify biological processes associated with differentially expressed genes (at p-adjusted < 0.01) between groups (Online Supp Table 1)

### miRNA prediction and analyses

Novel miRNAs were predicted using tools running in a local Galaxy environment (The Galaxy Community, 2022 update) and custom scripts. In a first step the FASTQ output from miRNA sequencing was combined and all reads were trimmed to remove adapter sequence using cutadapt with standard settings (*72*). The resulting insert sequences with sizes between 18 and 30 nucleotides were then subjected to the tools of the miRDeep2 package (*73*), and mapped to the Cobs3.1 genome using miRDeep2 mapper. Together with the genome these mappings served as input for the miRDeep2 miRNA prediction using standard parameters. For refinement of the miRNA prediction a set of mature miRNAs from relative species derived from miRBase (www.mirbase.org; version 22.1) was added to the miRNA prediction tool as well. As relative species were considered: *Apis mellifera*, *Dinoponera quadriceps*, *Nasonia longicornis* and *Polistes canadensis*. Because the names of the predicted miRNAs in miRDeep2 output follow the outdated mature/star scheme they were converted to the new 3p/5p name scheme of miRBase version 18 and later. In a second step the insert sequences of each of the separate libraries were counted against the set of predicted novel miRNAs using the miRDeep2 quantifier tool to get miRNA profiles for each of the 15 miRNA libraries (3 castes with each 5 replicates). The differential expression analysis between the 3 groups was performed using R package DEseq2 with alpha=0.05 (*74*).

## Acknowledgments

J.O. thanks Laurent Keller and Jan Medenbach for valuable comments. We also thank Helena Lowack for help with wet lab work and Andrea Bleckmann for help with confocal imaging. RNA sequencing was conducted at the NGS Core of the Leibniz Institute for Immunotherapy (LIT, University Regensburg and University Medical Center Regensburg, Germany).

## Funding

J.O. was supported by the Deutsche Forschungsgemeinschaft (OE 549/3-1,2). No funding sources were involved in study design, data collection and interpretation, or the decision to submit the work for publication.

## Author Contributions

J.O. and E.S. conceptualized the study. T.W., B.D., N.E., C.G., M.E., J.O. and E.S. produced the data and performed analyses. J.O. and E.S. wrote the manuscript, all authors contributed to the final version.

## Competing interests

Authors declare no competing interests.

## Data and materials availability

All data is available in the supplementary materials.

## List of supplementary materials

### Supplementary Figures

Fig S1 Embryogenesis in the ant *C. obscurior*.

Fig S2 Model of queen embryogenesis in the ant *C. obscurior*.

Fig S3 Expression of *dsx* in male, queen and worker embryos

Fig S4 A nest of *C. obscurior*

### Supplementary Tables

SuppTable1_mRNA_output.xlsx

SuppTable2_miRNA_output_Tables_Cobs3.1.xlsx

SuppTable3_FISHProbes.xlsx

SuppTable4mRNA.all.metrics.Limma.gold.xlsx

## Notes

### Competing Interest Statement

The authors have declared no competing interest.

### Summary of Updates

In response to critique by a colleague, we have removed the original Figs 1c and 1d and Supp Figs S2 and S4 for their suboptimal quality. The reasoning and conclusions are not affected by this.

