## Supplementary Figures for "Darwin’s “neuters” and the evolution of the sex continuum in a superorganism"

**Fig S1 Embryogenesis of the ant *C. obscurior*.**
**Fig S2 Model of queen embryogenesis in the ant *C. obscurior*.**
**Fig S3 Gene expression of *dsx* in male, queen and worker embryos.**
**Fig S4 A nest of *C. obscurior.***

**Fig S1 Embryogenesis of the ant *C. obscurior*.**

In *Cardiocondyla obscurior* embryonic development lasts for approximately nine days after egg laying (AEL). Strobl (Strobl et al., 2015) describe five embryonic stages in the red flour beetle *Tribolium castaneum*, which serve as a basis for a first description of the embryogenesis in this ant.

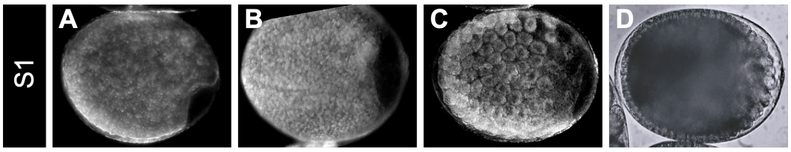


The amniotic fold is located at the posterior end of the embryo. (D) By the end of stage one the syncytial blastoderm has formed.

**Stage 1 (1 - 2 days AEL).** Characterized by the location of the amniotic fold at the posterior end of the embryo (A-C), followed by the formation of the syncytial blastoderm. At the posterior end of the syncytial blastoderm a region of the blastoderm thickens and forms the germ anlage, which later gives rise to the germ band (D).

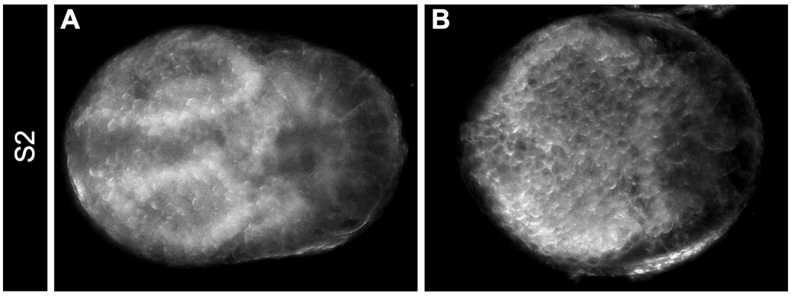


Ventral view at early gastrulation. The horseshoe-shaped amniotic fold is visible (View: anterior left, posterior right).

**Stage 2 (3 - 4 days AEL).** Defined by the gastrulation of the germ anlagen. Cells along the midline invaginate and proliferate (A, B). The single-layered germ anlage becomes two-layered. This process forms the germ band. Later, the anterior and posterior amniotic fold merge, giving rise to the serosa window. The serosa and amnion will be separated after the window closes ventrally. This step ends gastrulation (Chapman et al., 2013; Strobl et al., 2015).

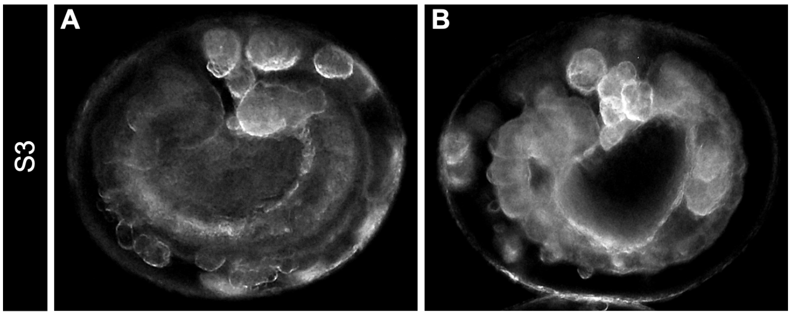


Lateral view of an embryo with fully elongated germ band. Large yolk cells nourish the developing embryo (View: anterior left).

**Stage 3 (5 - 6 days AEL).** The germ band is fully elongated and starts to retract dorsally. Separation of head and thorax is visible.

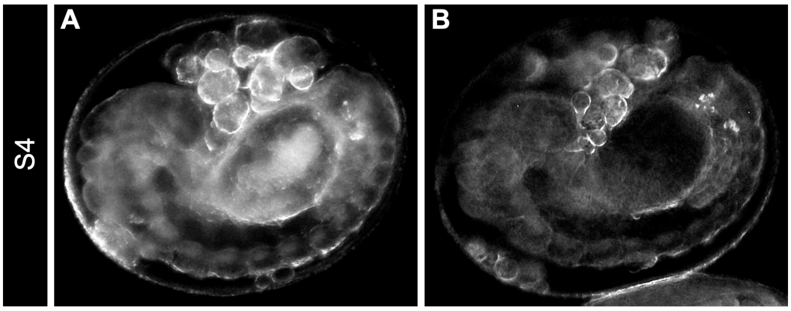


(A) Embryo with segmented germ band (gb). (B) Lateral view of an embryo with visible crystalline deposits (CD) (Schultner et al. 2022). md: mandibular segment; mx: maxillary segment; lb: labial segment

**Stage 4 (7 days AEL).** Characterised by the segmentation of the germ band and dorsal closure. The head, thorax and abdominal segments are well defined, making the gnathal segments (mandibular: md; maxilla: mx and labium: lb) distinguishable. Crystalline deposits are starting to be visible (B).

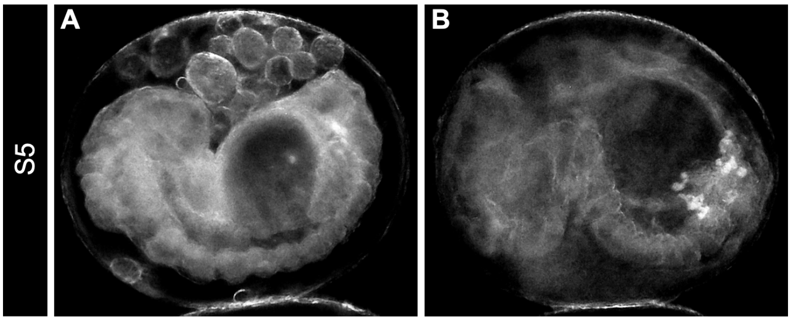


(A) Lateral view of a fully developed embryo. Segmentation of the germ band is completed. The head is in a ventral position. (B) Lateral view of an embryo with head turn to an anterior-ventral position. The crystalline deposits are visible as white dots

**Stage 5 (8 - 9 days AEL).** The head changes its orientation from a ventral to an anterior-ventral position. The embryo starts with muscular movement. Final step before the embryo hatches. The localization of crystalline deposits, likely comprising urate, is very distinct, making it possible to distinguish queen and worker embryos (B).

### Fig S2 Model of queen embryogenesis in the ant *C. obscurior*.

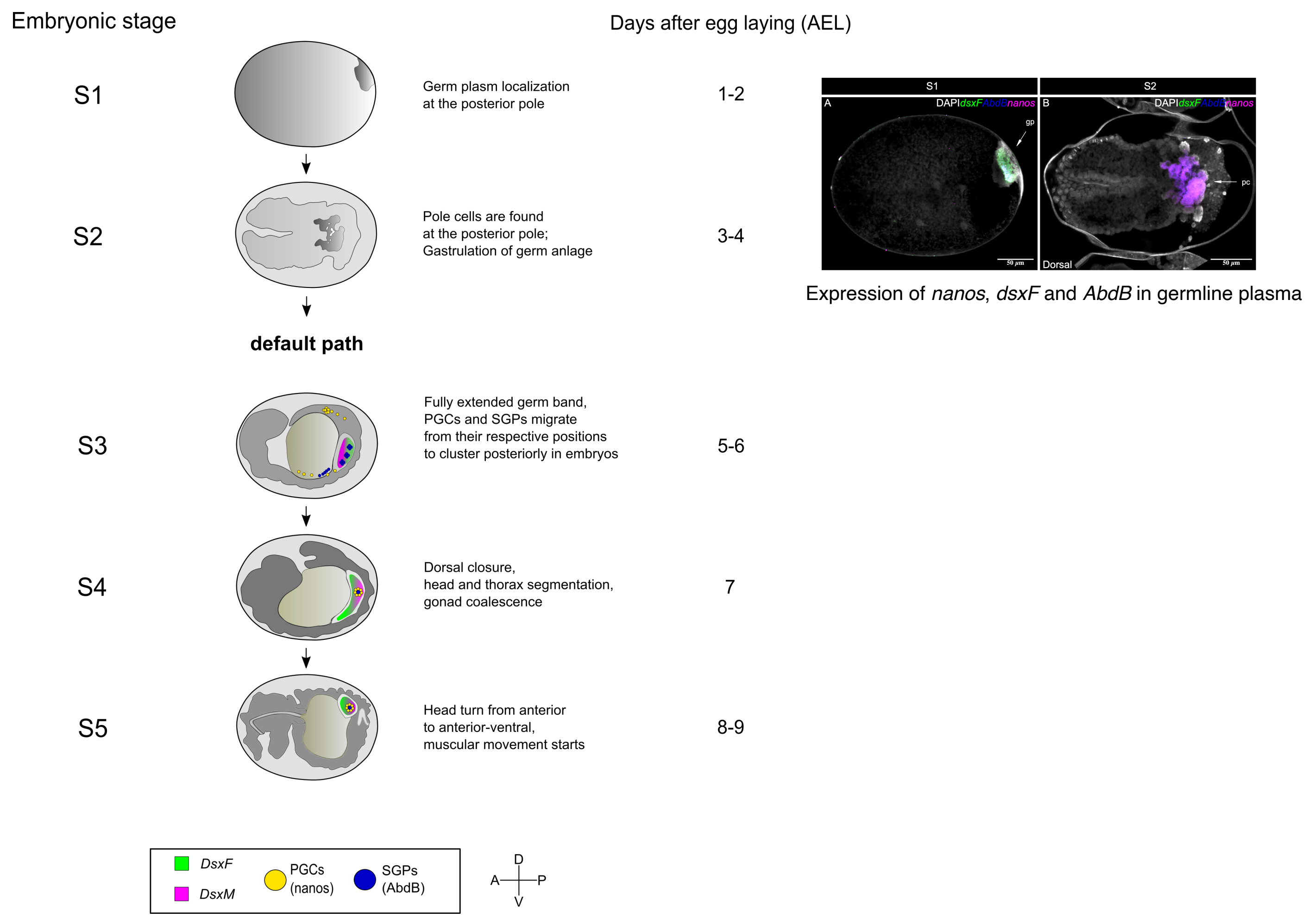

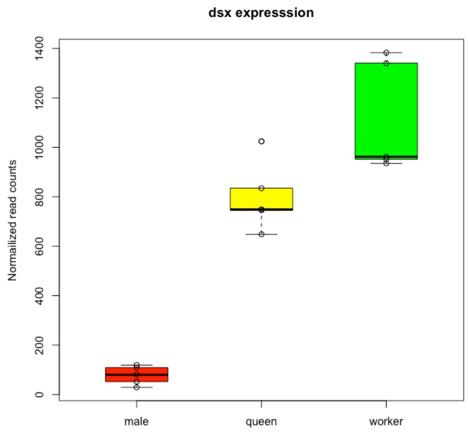


**Fig S3 Gene expression of *dsx* in male, queen and worker embryos.**

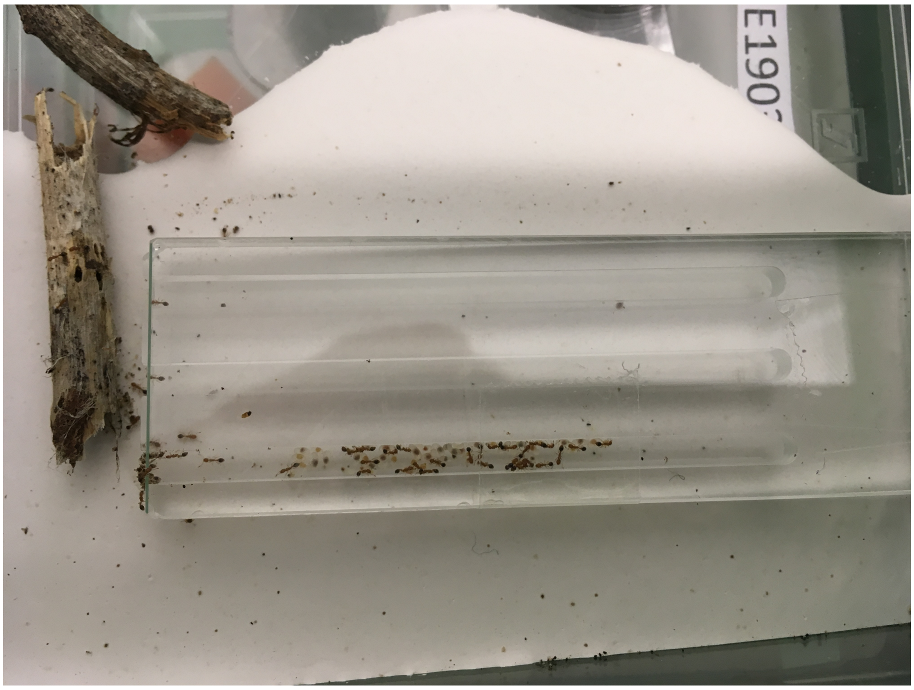

**Fig S4 A colony of *C. obscurior*.**

A freshly collected colony moves into its new home, a clean insert nest in a Sarstedt 100 mm x 100 mm x 20 mm petri dish. Some plaster-of-paris gives texture and helps maintain humidity. A queen is to the left, crouched at the entrance.
